# Synaptic pruning facilitates online Bayesian model selection

**DOI:** 10.1101/2024.05.15.593712

**Authors:** Ukyo T. Tazawa, Takuya Isomura

## Abstract

Identifying appropriate structures for generative or world models is essential for both biological organisms and machines. This work shows that synaptic pruning facilitates efficient statistical structure learning. We extend previously established canonical neural networks to derive a synaptic pruning scheme that is formally equivalent to an online Bayesian model selection. The proposed scheme, termed Bayesian synaptic model pruning (BSyMP), utilizes connectivity parameters to switch between the presence (ON) and absence (OFF) of synaptic connections. Mathematical analyses reveal that these parameters converge to zero for uninformative connections, thus providing reliable and efficient model reduction. This enables the identification of a plausible structure for the environmental model, particularly when the environment is characterized by sparse likelihood and transition matrices. Through causal inference and rule learning simulations, we demonstrate that BSyMP achieves model reduction more efficiently than the conventional Bayesian model reduction scheme. These findings indicate that synaptic pruning could be a neuronal substrate underlying structure learning and generalizability in the brain.

## 1. Introduction

Characterizing brain intelligence is a significant challenge in the field of biology. Biological organisms recognize the external milieu by inferring underlying structures. Despite the importance of structure learning—the process of identifying an appropriate structure to represent the environment (Braun et al., 2010; Lynn & Bassett, 2020; Tenenbaum et al., 2011)— its neuronal mechanisms in the brain are not yet fully understood (Tervo et al., 2016). The structure of neural networks is shaped by the pruning of synaptic connections during development (Sakai, 2020), and several computational studies have explored the advantages of synaptic pruning for memory and learning (Chechik et al., 1998; Scholl et al., 2021; Thomas et al., 2015). However, analyzing the relationship between biologically grounded synaptic pruning and information-theoretically derived structure learning is still required.

The free-energy principle has been proposed in theoretical neuroscience to account for various brain functions, including perception, learning, and action, in terms of minimizing variational free energy (Friston, 2010; Friston et al., 2006). It models the brain as an agent that performs variational Bayesian inference of external states and parameters by minimizing the variational free energy. These inferences rest upon a generative or world model that describes how latent variables in the external milieu generate sensory inputs. Recent works have showed that the dynamics of canonical neural networks that have a certain biological plausibility can be conceptualized as variational Bayesian inference under implicit generative models characterized by network structures (Isomura et al., 2022; Isomura & Friston, 2020). This lends the explainability of Bayesian inference to investigate the functional properties of biologically plausible neural networks. Additionally, Bayesian inference can be employed in selecting model structures (Friston et al., 2019; Friston et al., 2017; Jafarian et al., 2019). However, the neural substrate of Bayesian model selection is yet to be established.

Herein, we show that biologically plausible synaptic pruning enables structure learning in a Bayes-optimal manner by extending the previously established equivalence between neural networks and Bayesian inference. We reveal that incorporating synaptic pruning into a canonical neural network is equivalent to online Bayesian model selection, a method we term Bayesian synaptic model pruning (BSyMP). BSyMP adopts a set of connectivity parameters that enables switching between the presence (ON) and absence (OFF) of the model parameters. These connectivity parameters, in addition to states and other parameters, are updated by minimizing the variational free energy that is equivalent to a cost function for canonical neural networks. Thereby, BSyMP optimizes model structures via variational Bayesian inference of the hyper- or connectivity parameters of the generative model. Mathematical analyses show that BSyMP is guaranteed to prune parameters unnecessary for propagating hidden state information, resulting in efficient model reduction.

We compared the performance of BSyMP with that of the previously proposed Bayesian model reduction (BMR). BMR reduces the number of parameters in a learned full model by minimizing the variational free energy in a post hoc manner (Friston et al., 2019). Owing to online and continuous (i.e., gradual) pruning, BSyMP outperformed BMR in numerical simulations of causal inference and rule learning. BSyMP could deliver successful results even when the full model and BMR failed. Finally, we discuss possible neuronal substrates that realize synaptic pruning.

## 2. Results

### 2.1 Derivation of BSyMP from synaptic pruning

Herein, we consider a recurrent network of *N*_*x*_ neurons that receives *N*_*o*_-dimensional binary observations (sensory inputs). The activity of the neurons follows that of the previously defined canonical neural network model (Isomura et al., 2022). However, we extend the model by adding connectivity parameters (*C*_*W*_ and *C*_*K*_) that express the presence (ON) or absence (OFF) of synaptic connections, which is formulated as follows:

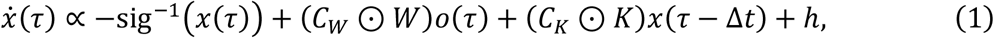

where 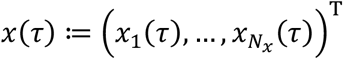 denotes the firing rates of the neurons; 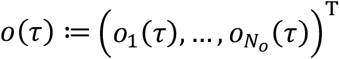 denotes binary observations; sig^-l^(*x*(*τ*)) denotes the inverse function of the sigmoid function of *x*(*τ*) that models the leakage current; 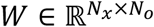 and 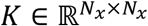 are the feedforward and recurrent synaptic strength matrices, respectively; the symbol ⊙ denotes the element-wise (Hadamard) product; Δ*t* > 0 is the delay in the recurrent input; and *h* represents the adaptive firing thresholds that depend on the synaptic strengths. The additional parameter matrices 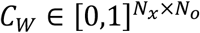 and 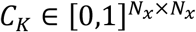 express the probability of the presence of synaptic connections, providing a major difference from the previous work. As shown in Eq. (1), when a certain element in *C*_*W*_ takes the value of 0, the signal from the corresponding synapse does not affect the neural activity. With this property, switching the values of *C*_*W*_ and *C*_*K*_ models the synaptic pruning process.

A previous work established a scheme for reverse-engineering cost function *L* for canonical neural networks from neural activity equations and showed the formal equivalence between *L* and the variational free energy *F* under a partially observable Markov decision process (POMDP) model (Isomura et al., 2022). Following the scheme, we reconstruct the cost function *L* from Eq. (1) (Fig. 1, left) and identify the variational free energy *F* that formally corresponds to *L* (Fig. 1, right).

**Fig. 1.**
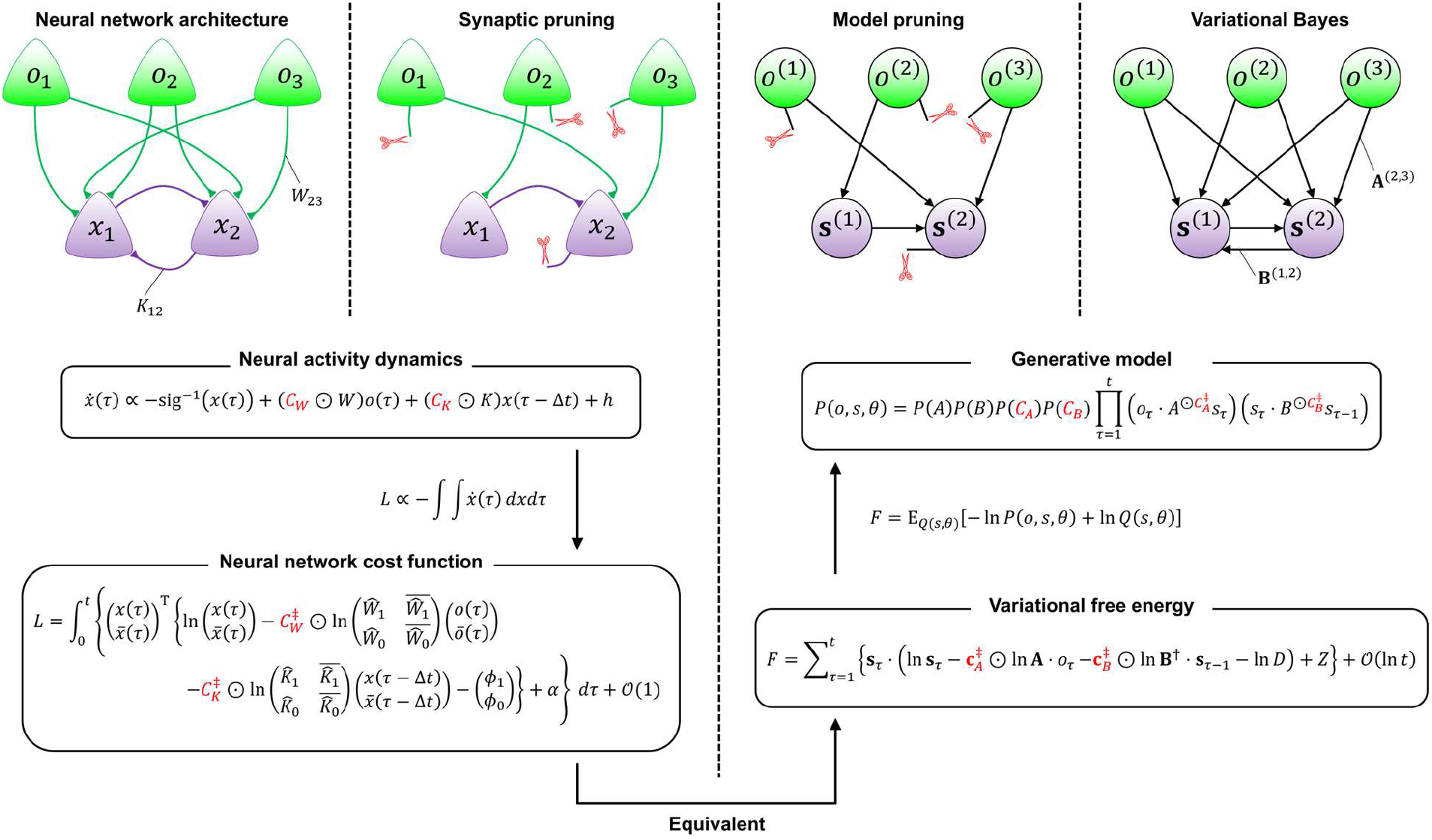
Equivalence between synaptic pruning in neural networks and Bayesian pruning of connections in a generative model. Top left: A recurrent neural network receives sensory inputs. The occurrence of synaptic pruning is mediated by the connectivity parameters *C*_*W*_ and *C*_*K*_. The neural activity dynamics are given by Eq. (1). Bottom left: The neural network cost function *L* is derived by integrating Eq. (1) with respect to *x* and time. The reconstructed *L* involves a free parameter *α* (i.e., integration constant). Bottom right: The variational free energy *F* equivalent to *L* is identified by extending the previously established equivalence between *F* and *L*. All variables are introduced in the main text and Table 1. Top right: The generative model *P*(*o, s, θ*) is reverse engineered from *F*. This enables the identification of the normalization factor *Z* that is involved in *F*. The details of the notations and derivations are provided in Section 4.1.

Subsequently, a biologically plausible generative model under which the neural network in Eq. (1) operates can be reverse engineered, as follows:

**Table 1.** Glossary of variables.

|  | Variables in neural networks | Variables in Bayesian inference |  |
| --- | --- | --- | --- |
| Sensory inputs | $o(\tau) := \left(o_1(\tau), \dots, o_{N_o}(\tau)\right)^T$<br>$\in \{0,1\}^{2 \times N_o}$ | $o_\tau := \left(o_\tau^{(1)}, \dots, o_\tau^{(N_o)}\right)^T$<br>$\in \{0,1\}^{2 \times N_o}$ | Observations |
| Neural activity | $x(\tau) := \left(x_1(\tau), \dots, x_{N_x}(\tau)\right)^T$<br>$\in [0,1]^{N_x}$ | $\mathbf{s}_\tau := \left(\mathbf{s}_\tau^{(1)}, \dots, \mathbf{s}_\tau^{(N_s)}\right)^T$<br>$\in [0,1]^{2 \times N_s}$ | State posterior |
| Feedforward synaptic strengths | $W \in \mathbb{R}^{N_x \times N_o}$ | $\mathbf{a} \in (\mathbb{R}^{2 \times 2})^{N_o \times N_s}$ | Likelihood mapping posterior |
| Recurrent synaptic strengths | $K \in \mathbb{R}^{N_x \times N_x}$ | $\mathbf{b} \in (\mathbb{R}^{2 \times 2})^{N_s \times N_s}$ | State transition posterior |
| Survival probability of feedforward synapses | $C_W \in [0,1]^{N_x \times N_o}$ | $\mathbf{c}_A \in [0,1]^{N_o \times N_s}$ | Connectivity posterior for $A$ |
| Survival probability of recurrent synapses | $C_K \in [0,1]^{N_x \times N_x}$ | $\mathbf{c}_B \in [0,1]^{N_s \times N_s}$ | Connectivity posterior for $B$ |
| Adaptive firing thresholds | $h \in \mathbb{R}^{N_x}$ | $D \in [0,1]^{2 \times N_s}$ | State prior (constant) |

Note: The variables in bold for the Dirichlet distributions (i.e., $\mathbf{a}$ and $\mathbf{b}$ ) indicate the concentration parameters, and those for categorical distributions (i.e., $\mathbf{c}_A$ , $\mathbf{c}_B$ , $\mathbf{s}_\tau$ ) indicate the posterior expectations of the corresponding variables in italics.

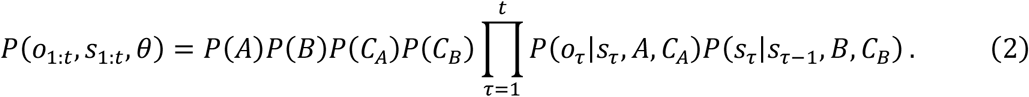

Here and throughout, *θ* ≔ {*A, B, C*_*A*_, *C*_*B*_} denotes a set of model parameters.

Equation (2) presents a POMDP model that generates *N_*o*_* observations 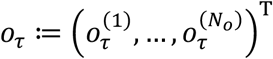 from *N_s_* hidden states 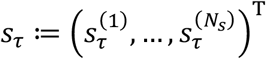 through the likelihood mapping *A* and its connectivity *C*_*A*_ in the form of a categorical distribution (also called a generalized Bernoulli distribution), 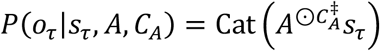 (Bishop, 2006). Each state and observation has a binary value. The hidden states *s*_τ_ depend on the previous states *s*_τ-l_, following the state transition matrix *B* and its connectivity 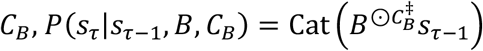. The prior belief about initial states is described by the block vector *D, P*(*s*_l_) = Cat(*D*), and throughout this work, *D* is a constant for simplicity. We consider the case in which *A* and *B* are factorized into products of 2 × 2 matrices *A*^(i,j)^ and *B*^(i,j)^. The new parameters 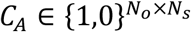 and 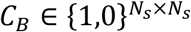 express the connectivity structure of graphical models. The double dagger ‡ on *C* denotes the block matrix of *C*, 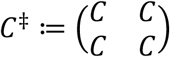, and the superscript ⊙ on *A* and *B* indicates the element-wise power. Owing to the factorial property of *A*, 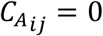 means the reduction (i.e., elimination or pruning) of *A*^(i,j)^ from the generative model, and the same applies to *B* (see Section 4.1 for further details). If 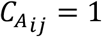 holds for all connectivities, Eq. (2) is reduced to a conventional full model without model reduction. The prior distributions of *A, B* and *C*_*A*_, *C*_*B*_ follow the Dirichlet distributions *P*(*A*) = Dir(*a*), *P*(*B*) = Dir(*b*) and categorical distributions *P*(*C*_*A*_) = Cat(*c*_*A*_), *P*(*C*_*B*_) = Cat(*c*_*B*_), respectively (Bishop, 2006). A simplified notation for the factorial structure was adopted here; please refer to previous works (Isomura et al., 2022; Isomura & Friston, 2020) for the detailed definition of a generative model that explicitly expresses the factorial structure.

Variational Bayesian inference is the process of optimizing an approximate posterior belief Q(*s*_1:t_, *θ*) based on a sequence of observations by minimizing the variational free energy as a tractable proxy for surprise minimization. Based on the mean-field approximation, the posterior distribution is expressed as follows:

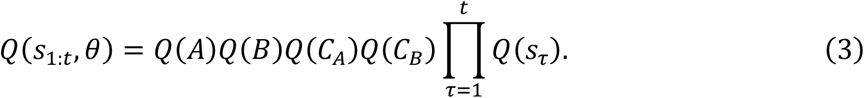

The marginal posteriors about *A, B, C*_*A*_, *C*_*B*_, *s*_*r*_ follow the Dirichlet distributions *Q*(*A*) = Dir(**a**), *Q*(*B*) = Dir(**b**) and categorical distributions *Q*(*C*_*A*_) = Cat(**c**_*A*_), *Q*(*C*_*B*_) = Cat(**c**_*B*_), *Q*(*s*_*r*_) = Cat(**s**_τ_), respectively. Here and throughout, the bold case variables (i.e., **c**_*A*_, **c**_*B*_, **s**_τ_, **A, B**) indicate the posterior expectations, while **a** and **b** denote the concentration parameters (Table 1). The analytical expression for the variational free energy under the abovementioned generative model and posterior beliefs is given as follows:

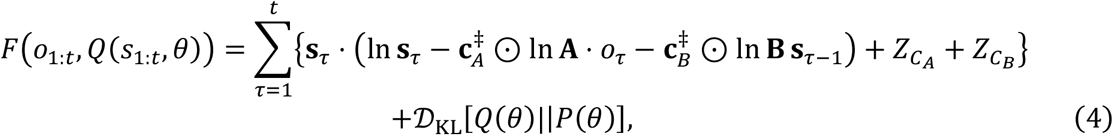

where 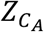 and 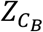 denote the normalization factors associated with **c**_*A*_ and **c**_*B*_, respectively; 𝒟_*KL*_[*Q*(*θ*)||*P*(*θ*)] denotes the complexity scored by the Kullback–Leibler divergence between *Q*(*θ*) and *P*(*θ*). Minimization of the variational free energy furnishes posterior beliefs. By solving the fixed point of the implicit gradient descent, *∂F*/*∂***c**_*A*_ = *0*, the update rule for 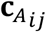 is formulated as follows:

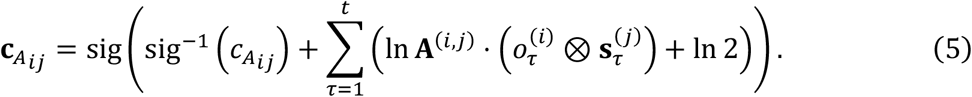

A similar equation holds for **c**_*B*_. The derivation details and explicit forms of the other posterior beliefs (i.e., **s**_τ_, **a**, and **b**) are provided in Section 4.2.

Equation (5) can be viewed as the dynamics of synaptic pruning in neural network formation. The probability of pruning is controlled by the similarities between the current synaptic strength (ln **A**^(i,j)^) and its change, as calculated by the Hebbian rule 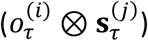 (Hebb, 1949). When these values are considerably different, their product becomes a large negative value over time, leading to the pruning of the corresponding connection. In addition, mathematical analyses show that an element of the connectivity parameter 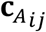 decreases and converges to 0 when the corresponding element of observation 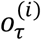 is randomly determined and independent of all elements of *s*_τ_, and the likelihood mapping posterior **A**^(i,j)^ and the mean state posterior 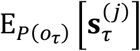 are sufficiently flat (refer to Section 4.3 for further details). This means that BSyMP is guaranteed to prune unnecessary connections in the generative model as long as the prior of the likelihood mapping and the mean state posterior are unbiased.

In short, the connectivity parameters *C*_*A*_ and *C*_*B*_ provide an implementation of the Bayesian model selection that enables switching between the presence (ON) and absence (OFF) of each element of matrices *A* and *B*. As shown in Fig. 1, this model reduction is formally equivalent to synaptic pruning in neural network formation, which is mediated by *C*_*W*_ and *C*_*K*_ in Eq. (1). Table 1 summarizes the correspondence between variables in neural networks and those in variational Bayes.

### 2.2. Numerical simulation 1: Blind 175 source separation

The performance of BSyMP was evaluated using a blind source separation (BSS) task, in which the performance was expected to improve with appropriate model reduction. This BSS task was analogous to a visual shape recognition task in which sources 1 and 2 displayed a low-contrast circle and rectangle, respectively, on a 32 × 32 pixel screen with a certain fidelity following a likelihood mapping (Fig. 2a). Each pixel on the display took either 0 or 1 stochastically, where some parts of the shapes overlapped. Thus, the agent needed to disentangle the input signals to perceive the hidden states. The field of view of the agent comprised 1024-dimensional observations, and the states of the two sources (ON/OFF) changed randomly. This setup was effectively expressed as a generative model without matrix *B*; thus, the agents optimized only **s**_τ_, **a**, and **c**_*A*_.

**Fig. 2.**
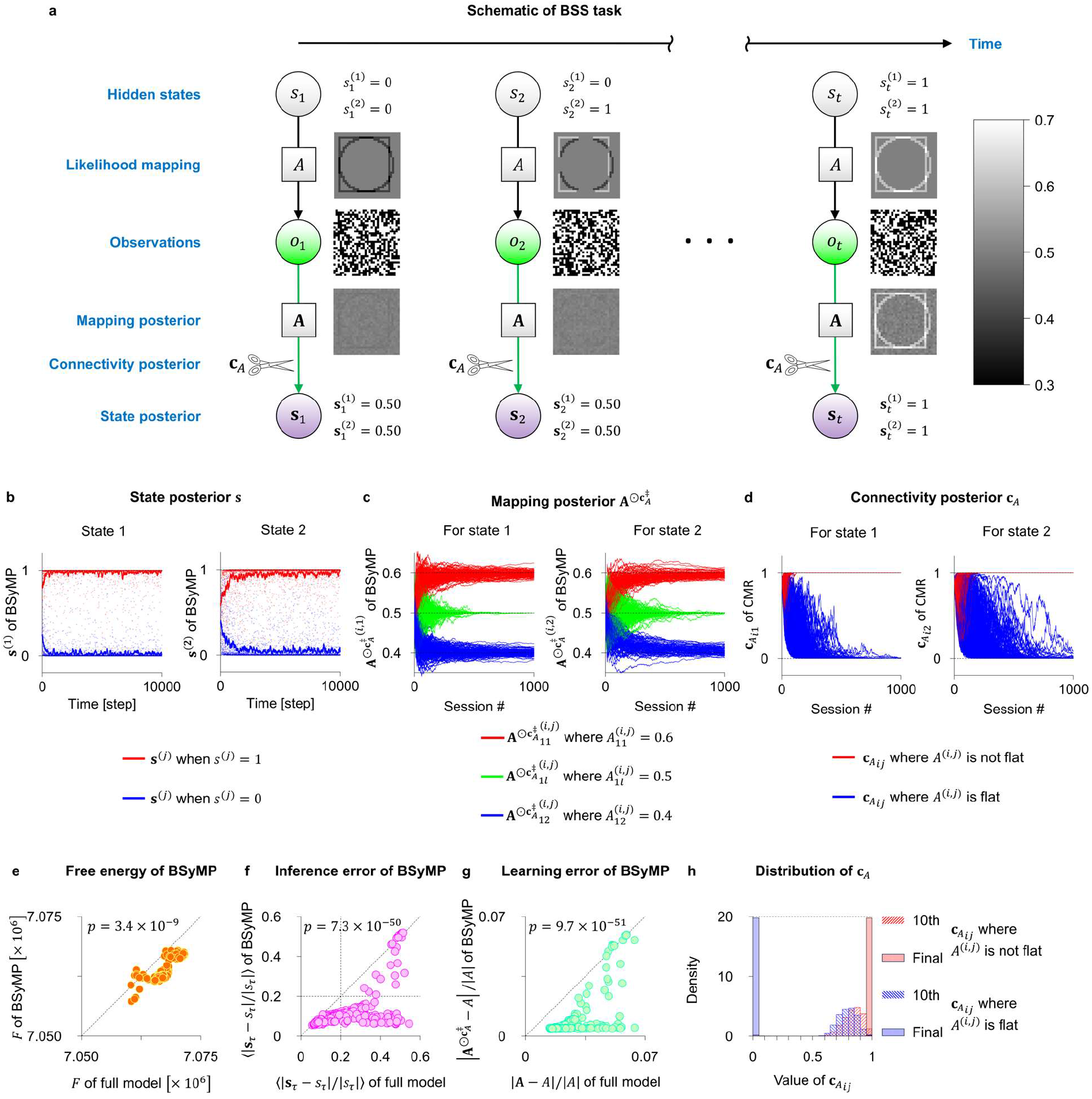
Performance of BSyMP agent on BSS task. (**a**) A schematic of the task, illustrating a generative process with one likelihood mapping out of three. Two hidden sources *s*_*r*_ randomly take the value of 1 (ON) or 0 (OFF). Binary observations *o*_*r*_ are stochastically generated from *s*_*r*_ through the likelihood mapping *A*, *P*(*o*_*τ*_|*S*_*τ*_, *A*) = Cat(*As_τ_*). The three vertically aligned images depict *As_τ_*, *o*_τ_, and *As_τ_* in order from top to bottom. (**b**) The posterior beliefs about two states during a single simulation. The red and blue dots indicate 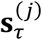 when the true hidden state 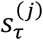 is 1 or 0, respectively. The red and blue trajectories indicate the moving averages of the red and blue dots with a window length of 100 time points. (**c**) The posterior beliefs about the likelihood mapping during a single simulation. The elements in the posterior beliefs are grouped and colored according to the value of the corresponding elements in the true mapping, that is, 0.6 (red), 0.4 (blue), or 0.5 (green). (**d**) The posterior beliefs about the connectivity during a single simulation. The elements in the posterior beliefs are grouped and colored according to whether the corresponding submatrix in the true mapping is flat (blue; to be reduced) or not (red; to be retained). (**e**) A comparison of the free energy between BSyMP and the full model. In (**e**), (**f**), and (**g**), each point represents data obtained in a single simulation. Throughout this work, the *p* value was obtained by the two-sided Wilcoxon signed-rank test for the pair. (**f**) A comparison of the state inference error between BSyMP and the full model. The error was evaluated by the mean of the Euclidean norm ratio |**s**_τ_ − *s*_τ_|/|*s*_τ_| in the test session. (**g**) A comparison of the parameter learning error between BSyMP and the full model. The error was evaluated by the Frobenius norm ratio 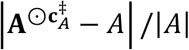 for BSyMP and |**A** − *A*|/|*A*| for the full model at the end of the training sessions. (**h**) The distribution of the posterior beliefs about the connectivity. All the data at the 10th and final training sessions of 300 simulations were counted in 0.05 intervals. Please refer to Section 4.4 for further details.

Three agent types (full model, BMR, and BSyMP) were trained with identical input sequences. The agents optimized **s**_*T*_ at every time point; conversely, they optimized **a** and **c**_*A*_ at every 10 time points, which we refer to as a training session. One simulation comprised 1000 training sessions and a subsequent test session. In every test session, the agents inferred **s**_τ_ for 1000 time-point-length test data that were newly generated from the same likelihood mapping used in the training sessions. We prepared three likelihood mappings, generating inputs that differed in the size of the circle and rectangle and the overlap between them. The simulation was repeated 100 times for each mapping (i.e., 300 simulations in total).

The agents with BSyMP successfully self-organized to infer the hidden sources as a result of successful learning of posterior beliefs about the likelihood mapping (Figs. 2b and 2c). The result was underwritten by an appropriate model reduction, as depicted by the efficient reduction of the redundant connections (Fig. 2d). When the source and observation were independent of one another, BSyMP reliably reduced the connectivity between them to 0 (Fig. 2d; blue lines). This result was consistent with the mathematical analysis described in the previous subsection and Section 4.3. Figure 2d further shows that the connectivity was retained if the source and the observation were not independent (red lines), indicating that no over-pruning occurred in this task. Owing to these desirable properties of the connectivity parameters, the variational free energy of the BSyMP agent was smaller than that of the full model agent (Fig. 2e), and the inference and learning performance of the BSyMP agent in this BSS task significantly exceeded those of the full model agent (Figs. 2f-h). For ease of discussion, we labeled a simulation as successful if the state inference error in the test session was less than 20%, and as a failure otherwise (Fig. 2f; vertical and horizontal dashed lines). Under this criterion, the success rate of the full model agent was 45.0% (135/300), whereas that of the BSyMP agent was 92.3% (277/300). These results demonstrate that BSyMP achieved appropriate model reduction and significant effects on the performance of the BSS task.

Subsequently, BSyMP was compared with a standard Bayesian model reduction method, referred to as BMR. BMR compares the free energy of the learned full model with that of the reduced version of the model at a small computational cost. Thus, BMR is a post hoc model reduction method that is based on an already acquired model.

Although the BMR agent decreased the variational free energy more than the BSyMP agent, the performance of the BMR agent was close to that of the full model and did not exceed that of the BSyMP agent (Figs. 3a-c). The success rate of BMR was 46.7% (140/300), which was only 1.7% higher than that of the full model, whereas the rate of BSyMP was twice as high as that of the full model. The inconsistency between the free energy and the performance was arguably because BMR decreased the complexity term 𝒟_*KL*_[*Q*(*A*)||*P*(*A*)] in the free energy by heuristically altering priors (Friston et al., 2017), but this free energy reduction had little contribution to the performance in this setting. By contrast, BSyMP could prune synapses by evaluating the likelihood through its construction. Consistent with the performance, the accuracy of model reduction by BMR is worse than BSyMP (Fig. 3d).

**Fig. 3.**
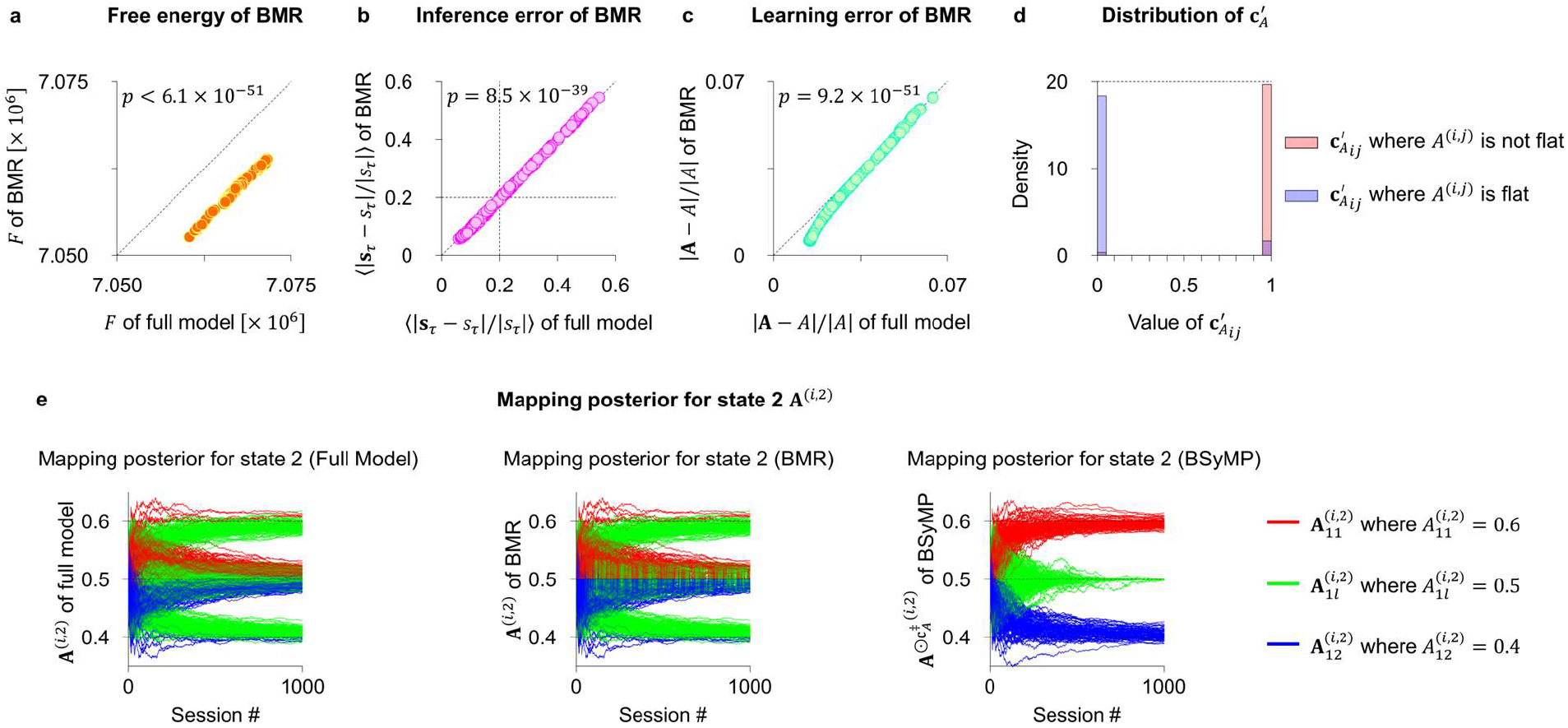
Comparison of BSyMP with BMR on BSS task. (**a**) A comparison of the free energy between BMR and the full model. (**b**) A comparison of the state inference error between BMR and the full model. (**c**) A comparison of the parameter learning error between BMR and the full model. (**d**) The distribution of the BMR’s counterpart, 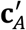, to the connectivity parameter in BSyMP, **c***_A_*, at the end of the training sessions. Here, 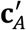 has the same number of elements as **c**_*A*_, and each element in 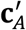 takes a binary value, indicating whether the corresponding submatrix of the mapping posterior is reduced (0) or not (1). (**e**) The posterior beliefs about the likelihood mapping regarding state 2 during a single simulation obtained by three methods. Note that the time course of the posterior beliefs of a BMR agent was calculated by performing BMR at every session.

We attribute the underperformance of BMR to the post hoc manner of its model reduction. This hypothesis was supported by the comparison of parameter trajectories. Figure 3e illustrates the typical outcomes of three approaches in a simulation. Owing to stochasticity in the generative processes and priors, the agent with the full model often (in 55% of the simulations) failed to learn the likelihood mapping (Fig. 3e, left panel). BMR could reduce the redundant elements but could not change the learning process because it operates in a post hoc manner (Fig. 3e, center panel). By contrast, as BSyMP adopts online reduction during learning, it could learn the likelihood mapping based on the state inference that was improved by the model reduction, resulting in its reliable success (Fig. 3e, right panel). While naive BMR operates in the post-hoc manner, BMR can be applied to operate online (Friston et al., 2017). However, the online version of BMR performed the same level as the post-hoc BMR, because of its heuristics (Appendix A). In summary, we demonstrated that BSyMP successfully eliminated connections independent of the sources and improved the performance of the BSS task more significantly than related methods.

### 2.3 Numerical simulation 2: Rule learning

The performance of BSyMP was further evaluated using a rule-learning task in which the agents learned the transition rule behind the visual inputs in addition to BSS. The agents received observations comprising 32 × 96 pixels, which is analogous to three screens. Four sources on each screen stochastically displayed a circle, rectangle, x, and plus, respectively (Fig. 4a). In other words, the agents received 3072 binary observations that depend on the states of the 12 sources (ON/OFF). The states changed according to the transition rule described by matrix *B*. The agents had to learn the rule to predict the next state of the sources.

**Fig. 4.**
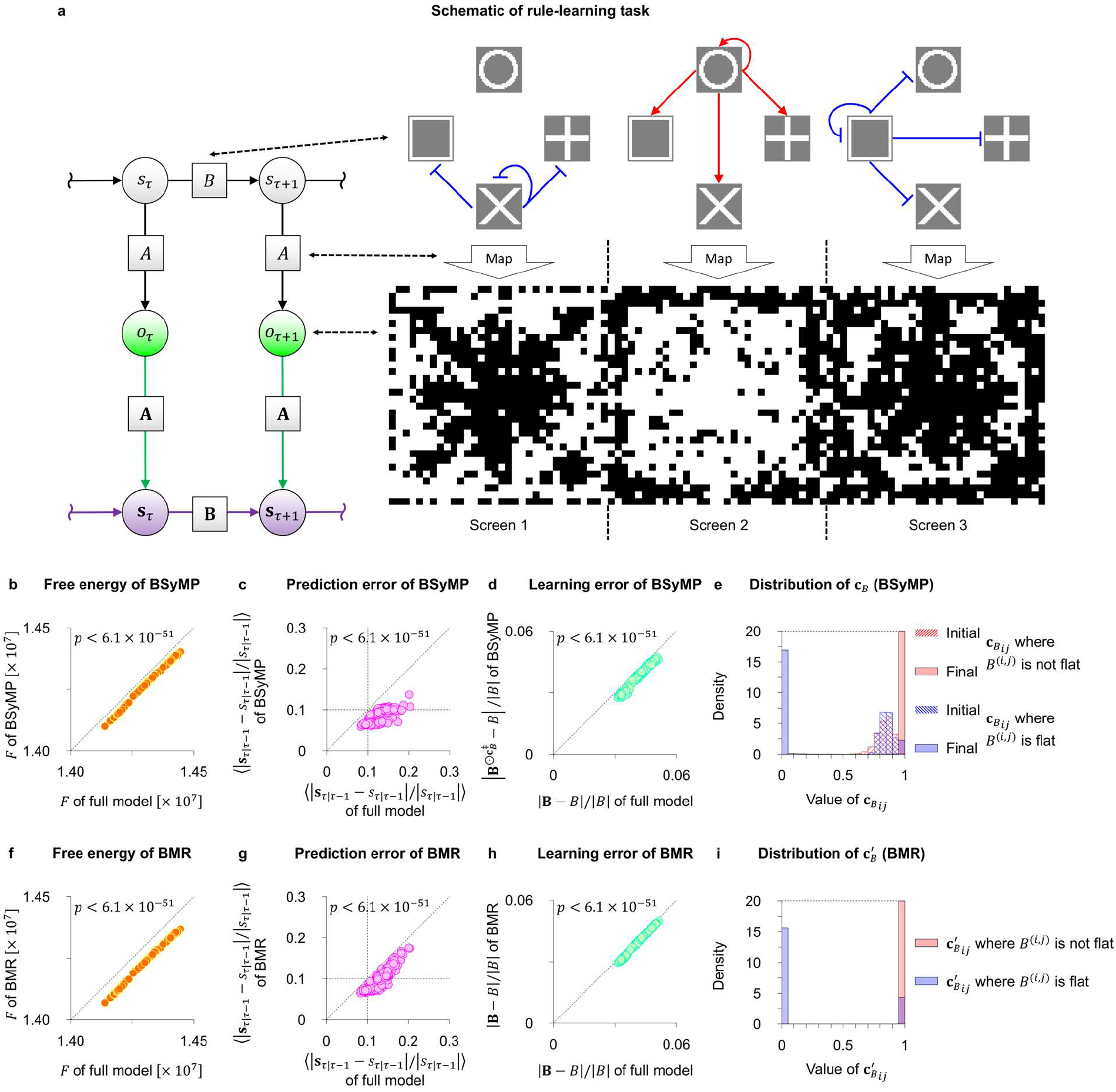
Performance of BSyMP agent on rule-learning task. (**a**) A schematic of the task. The factor graph on the left side illustrates a generative process. Images on the right side depict one of three transition rules and one example of observations at a moment. The states of the sources stochastically change depending on the states at the previous time point. The red arrows from one source *s*^(*j*)^ to another source *s*^(*i*)^ indicate the existence of the positive connection from 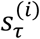 to 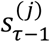, whereas the blue T-shaped lines indicate the negative connection. Each source displays a shape in one of three 32 × 32-pixel screens and has no effect on the other 32 × 64 pixels. The agent optimizes the posterior beliefs about *s*_*r*_, *A, B, C*_*A*_, and *C*_*B*_. (**b**) A comparison of the free energy between BSyMP and the full model. (**c**) A comparison of the state prediction error between BSyMP and the full model. The error was evaluated by the mean of the normalized Euclidean distance between the actual state and agent’s prediction |**s**_τ|τ-l_ − *s*_τ| τ-l_|/ |*s*_τ| τ-l_| in the test session. Please refer to Section 4.5 for the definitions of **s**_τ| τ-l_ and *s*_τ| τ-l_. (**d**) A comparison of the tule learning error between BSyMP and the full model. The error was evaluated by the Frobenius norm ratio 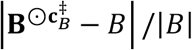 for BSyMP and |**B** − *B*|/|*B*| for the full model at the end of the training sessions. (**e**) The distribution of the posterior beliefs about the connectivity parameter. (**f**) A comparison of the free energy between BMR and the full model. (**g**) A comparison of the state prediction error between BMR and the full model. (**h**) A comparison of the rule-learning error between BMR and the full model. (**i**) The distribution of the BMR’s counterpart, 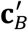, to the connectivity parameter in BSyMP, **c**_*B*_, at the end of the training sessions.

We trained the three agent types in the same manner as in the BSS task. The agents optimized **s**_τ_ at every time point and optimized **a, b, c**_*A*_, and **c**_*B*_ at every 100 time points during the training sessions. In this case, we provided the agents with accurate and strong prior beliefs about the likelihood mapping *A* to focus on the learning in matrix *B*, thereby ensuring accurate state inference after obtaining the observations. One simulation comprised 100 training sessions and a subsequent 1000 time-point-length test session in which the prediction error for the upcoming states was evaluated. We prepared three transition rules, and the simulation was repeated 100 times for each rule (i.e., 300 simulations in total).

The BSyMP agents scored a smaller variational free energy than the full model agent and successfully predicted the upcoming hidden states by learning the posterior beliefs about the transition rules (Figs. 4b-d). The success was underwritten by the appropriate pruning of redundant recurrent connections between the state posteriors (Fig. 4e). The prediction and learning performance of the BSyMP agent in the rule-learning task significantly exceeded those of the full model agent (Figs. 4c and d) owing to the desirable properties of the connectivity parameters. For ease of discussion, we labeled a simulation as successful if the state prediction error in the test session was less than 10%, and as a failure otherwise (Fig. 4c). Under this criterion, the success rate of the full model agent was 9.0% (27/300), whereas that of the BSyMP agent was 82.3% (247/300). These results show that BSyMP achieved appropriate model reduction and significant effects on the performance of the rule-learning task.

Subsequently, BSyMP was compared with BMR. As in the BSS task, although BMR decreased the variational free energy more than BSyMP owing to the decrease in complexity (Fig. 4f), and the performance of the BMR agent exceeded that of the full model agent (Figs. 4g and h), the success rate of the BMR agent, 58.7% (176/300), was less than that of BSyMP. This performance difference could not be attributed to the difference in the manner of model reduction (i.e., online or post hoc) because the model reduction had no significant effect on learning matrix *B* in this rule-learning task, owing to accurate state inference, unlike in the BSS task. Nevertheless, pruning unnecessary connections is more challenging in this task than in the BSS task because some sources can be correlated with others despite the absence of direct connections. For example, in the rule shown in Fig. 4a, the state of the rectangle on screen 1 at time τ is negatively correlated with the state of the plus on screen 1 at time τ + 1. The blue bars in Figs. 4e and 4i show that BSyMP was better at pruning these “false positive connections” than BMR, which can explain the outperformance of BSyMP in rule-learning tasks. In summary, we demonstrated that BSyMP successfully eliminated unnecessary connections between the sources and improved the rule-learning performance to a greater extent than the previously developed method.

## 3. Discussion

We derived the biologically plausible synaptic pruning rule, BSyMP (Eq. (5)), from the cost function minimization based on the equivalence between the dynamics of the canonical neural networks and variational Bayesian inference. Equation (5) states that the probability of pruning depends on the similarities between the current synaptic strength (ln **A**^(*i*,*j*)^) and its change calculated by the Hebbian rule 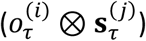 during the latest session. The Hebbian plasticity in the brain can be achieved by long-term potentiation (LTP) and depression (LTD), and its molecular mechanisms are well studied (Malenka & Bear, 2004). The Hebbian term in Eq. (5) can be encoded in the molecules in the postsynaptic neuron (Nicoll, 2017; Scannevin & Huganir, 2000), while the values of synaptic strengths lie in the density of AMPA receptors in the synaptic plasma membrane. Thus, it is possible that the molecular mechanisms in the postsynaptic neuron compute the correlation between ln **A**^(*i*,*j*)^ and 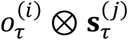 to determine which synapses are pruned. Experimentally observed similarities between LTP, LTD, and synaptic pruning—in their physiological properties and molecular mechanisms—further support the biological plausibility of our model, highlighting its potential to guide the experimental targets to shed light on the mechanisms underlying synaptic connectivity control (Helias et al., 2008; Le Bé & Markram, 2006; Lüscher et al., 2000).

One can contrast BSyMP with earlier pruning methods. Pruning connections with small weight is the most basic and widely used rule to prune artificial neural networks (Hagiwara, 1994; Han et al., 2015, 2016; Hoefler et al., 2021). Our analysis shows that BMR constitutes a family of weight-base pruning rules (please refer to Appendix B for details). In addition to weight-based methods, pruning methods based on the activity of neurons exist (Scholl et al., 2021; Sietsma & Dow, 1991; Sun et al., 2015). BSyMP utilizes temporal consistency to organize the network structure by comparing synaptic weight (i.e., the history of neural activity in the past) with neural activity in the latest session. This allows the neural network to spontaneously detect changes in the statistical features of neural activity. This would be useful for reorganizing the network structure flexibly to adapt to environmental changes, enabling BSyMP to outstand in the activity-based pruning rules. This might be a mechanistic explanation of the dynamics of the network structure in the brain (Ziv & Brenner, 2018). Moreover, most pruning methods, including BMR, prune synapses in a binary manner, and their processes involve some heuristics. BSyMP, by contrast, computes the probability of the presence of synapses and prunes synapses in a continuous (i.e., differentiable) manner, thereby enabling stochastic gradient descent. This rule is formally derived from variational free energy minimization, achieving more accurate pruning than the binary methods. This is a great advantage over heuristic methods. Appendix A delves into this contrast analytically.

Biological organisms utilize structural learning to identify the structure of their environment (Braun et al., 2010; Lynn & Bassett, 2020; Tenenbaum et al., 2011). Structure learning facilitates generalization by transferring the structures obtained in the past to newly encountered environments (Collins & Frank, 2013). Our work suggests that synaptic pruning plays a vital role in structure learning in the brain. This would be useful for designing next-generation artificial intelligence with high data efficiency and robustness and for understanding the mechanisms underlying the generalizability of the brain and its attenuation caused by psychiatric disorders (Kasai et al., 2021).

## Conclusion

In this work, we derived a reliable online Bayesian model selection scheme, BSyMP, that can be implemented by canonical neural networks equipped with synaptic pruning. Mathematical analyses revealed that BSyMP is guaranteed to prune unnecessary connections in the generative model. Numerical simulations of BSS and rule-learning tasks demonstrated the advantage of BSyMP over the conventional BMR scheme. These findings suggest that synaptic pruning can be a possible neuronal substrate underlying structure learning and generalizability in the brain.

## 4 Methods

### 4.1 Derivation of generative model from neural network

The generative model of BSyMP originates from the canonical neural network described in Eq. (1). We derive the cost function *L* for canonical neural networks from Eq. (1) and identify the variational free energy *F* under a certain POMDP generative model that formally corresponds to *L*. For this purpose, we introduce excitatory and inhibitory synapses for feedforward circuits *W*_1_ and *W*_*O*_ that satisfy *W* = *W*_1_ − *W*_*O*_ and those for recurrent circuits *K*_1_ and *K*_*O*_ that satisfy *K* = *K*_1_ − *K*_*O*_. The corresponding adaptive firing thresholds are denoted as *h*_1_ and *h*_*O*_, which are functions of *W*_1_, *K*_1_, *W*_*O*_, *K*_*O*_, *C*_*W*_, *C*_*K*_ and satisfy *h* = *h*_1_ − *h*_*O*_. This treatment separates the positive (variables with subscript 1) and negative (variables with subscript 0) effects on neural activity. As the gradient descent on *L* yields Eq. (1), *L* can be derived by integrating Eq. (1) with respect to neural activity *x* and time *τ*, as follows:

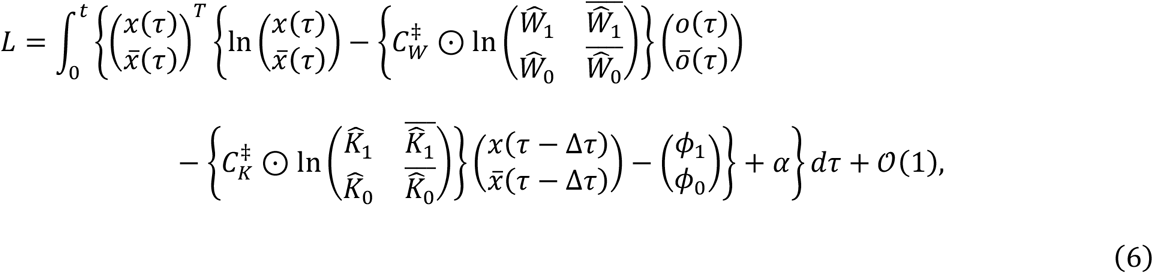

where 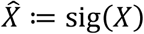 denotes the sigmoid function, 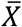 is defined as 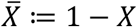 for any variable *X*, and *α* denotes the integration constant that is independent of *x, W*, and *K* but can depend on *C*_*W*_ and *C*_*K*_.

The previous work established the equivalence between the cost function of neural networks without connectivity parameters *C*_*W*_ and *C*_*K*_, denoted as *L*_*woC*_, and the variational free energy under the POMDP model *F*_*woC*_ (Isomura et al., 2022). When all elements of *C*_*W*_ and *C*_*K*_ are equal to 1, Eq. (6) is reduced to *L*_*woC*_ and the corresponding variational free energy is expressed as follows:

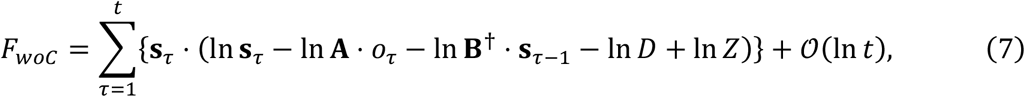

where **B**^†^ ≔ **B**^T^diag(*D*)^-1^ denotes the inverse transition matrix and *D* is the prior belief about the hidden states, in which ln **B s**_τ-1_ = ln **B**^†^ ⋅ **s**_τ-1_ + ln *D* holds. For simplicity, we consider *D* as a constant. The partition function *Z* is a constant that is independent of **s, A**, and **B**. Although *Z* was omitted in the previous work because its derivatives with respect to **s, A**, and **B** are 0, it is involved in the generative model as a normalization factor. This *Z* is a counterpart to *α* in *L*. With this equivalence, the variational free energy *F* that is equivalent to cost function *L* in Eq. (6) takes the following form:

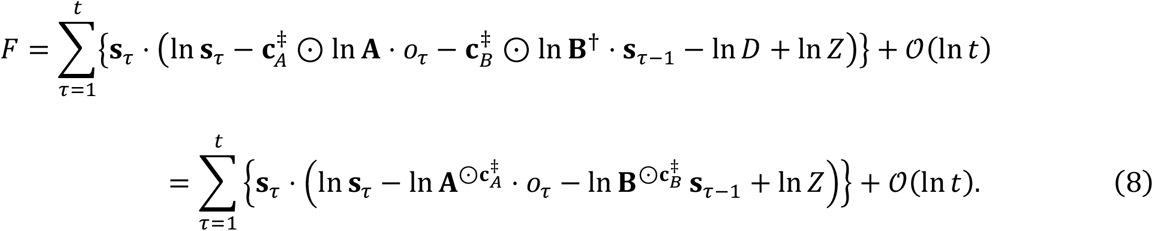

The new parameters **c**_*A*_ and **c**_*B*_ are the counterparts to *C*_*W*_ and *C*_*K*_ in Eq. (6), respectively. At this point, the POMDP generative model underlying Eq. (8) can be reverse engineered by adding the roles of **c**_*A*_ and **c**_*B*_ to the conventional POMDP, as per Eq. (2) in Section 2:

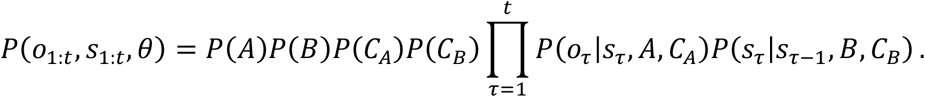

In the following subsection, we derive the explicit form of the free energy of BSyMP and confirm that *Z* in Eq. (8) is independent of **s, A**, and **B**, which is necessary for making this Bayesian formulation equivalent to canonical neural networks with synaptic pruning.

The explicit form of this generative model is expressed as follows:

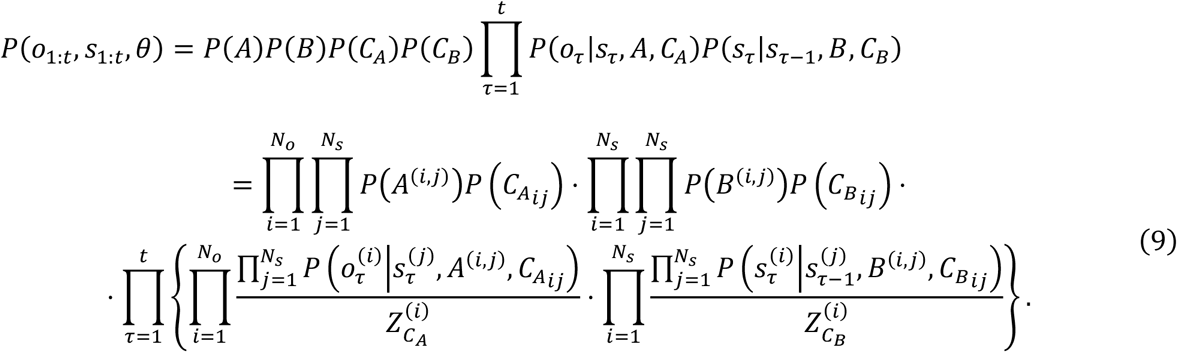

The prior beliefs about *A*^(*i*,*j*)^, *B*^(*i*,*j*)^ and 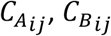 follow the Dirichlet distributions *P*(*A*^(i,j)^) = Dir(*a*^(i,j)^), *P*(*B*^(i,j)^) = Dir(*b*^(i,j)^) and categorical distributions 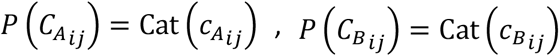, respectively. The initial states follow a categorical distribution: *P*(*s*_1_) = Cat(*D*). As *C*_*A*_ takes the value of 1 or 0, the following equation holds:

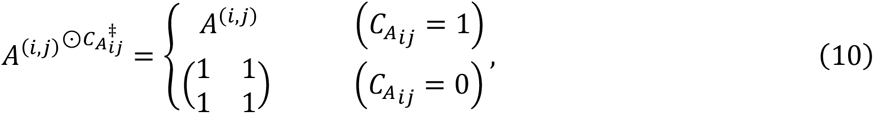

which means that 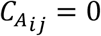 reduces *A*^(i,j)^ from the model owing to the factorial property of the model. The same applies to *B*. As mentioned in Section 2, 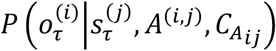 and 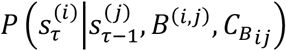 are given by categorical distributions 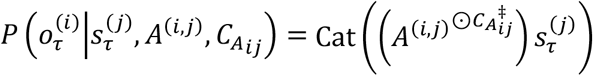 and 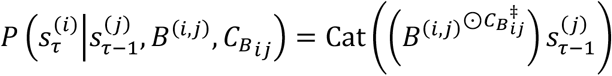 respectively.

The partition functions 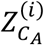 and 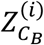 serve as normalization factors to counterbalance the deviations of 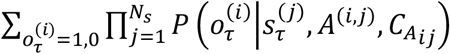 and 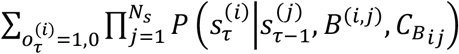 from 1, which are expressed as

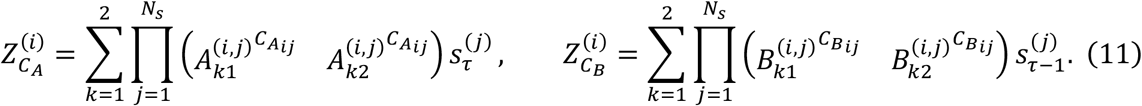

### 4.2 Variational free energy and belief update rule

In this subsection, we describe the explicit forms of posterior beliefs and variational free energy and derive belief update rules that minimize variational free energy. The explicit form of Eq. (3) (adopting the factorial structure) can be expressed as

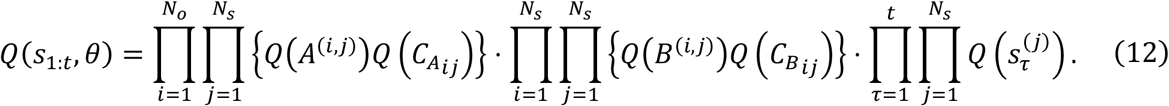

The marginal posterior beliefs about *A*^(i,j)^, *B*^(i,j)^, 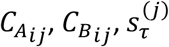 follow the Dirichlet distributions *Q*(*A*^(i,j)^) = Dir(***a***^(i,j)^), *Q*(*B*^(i,j)^) = Dir(***b***^(i,j)^)and categorical distributions 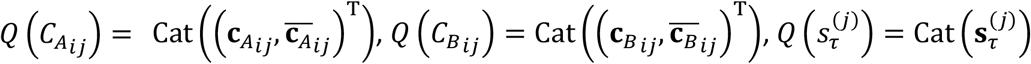, respectively.

Under the generative model defined in Eq. (9), the variational free energy is expressed as follows:

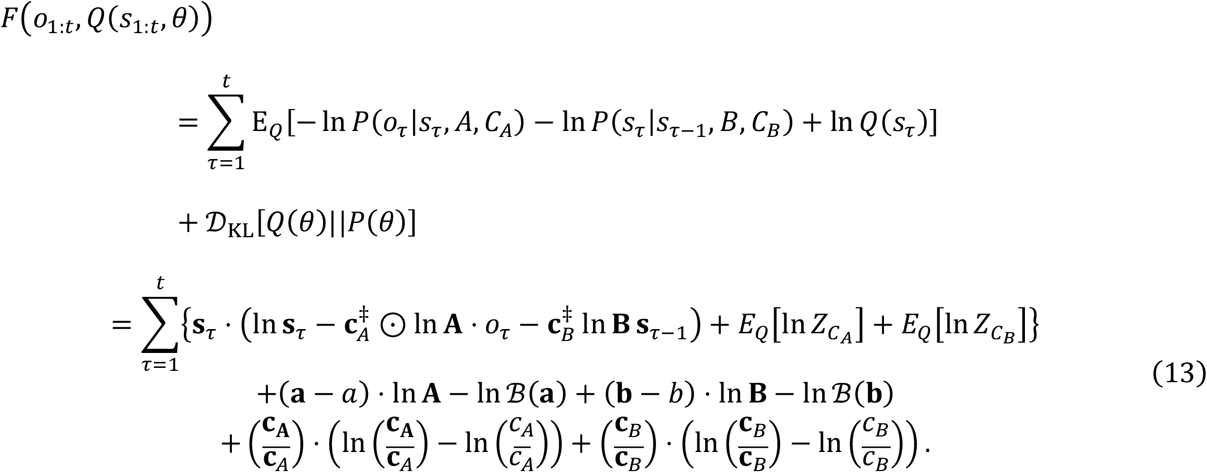

Note that factorial structures (i.e., summations over *N*_*s*_ and *N*_*o*_) are implicitly involved for simplification. In the above, 𝒟_*KL*_[*Q*(*θ*)||*P*(*θ*)] denotes the complexity scored by the Kullback– Leibler divergence between *Q*(*θ*) and *P*(*θ*), 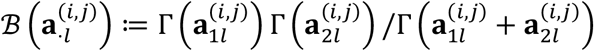 denotes the beta function characterized by the gamma function Γ(·), 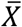 denotes 1 − *X*, and In 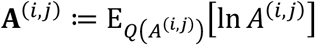 denotes the posterior expectation of ln *A*^(i,j)^. Using a first-order approximation on *A*^(i,j)^ and *B*^(i,j)^, E_*Q*_[ln *Z*] can be calculated as follows:

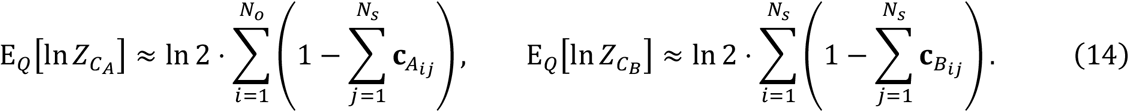

The detailed derivations of this approximation are provided in Appendix D. The term 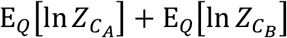 is independent of **s, A**, and **B** and corresponds to ln *Z* in Eq. (8), which also corresponds to *α* in Eq. (6) that is independent of *x*, *W*, and *K*. Thus, this relationship confirms the formal equivalence between the variational free energy under BSyMP (Eq. (8)) and the cost function for canonical neural networks with synaptic pruning (Eq. (6)).

Variational free energy minimization yields posterior beliefs about the hidden states and parameters. By solving the fixed point of the implicit gradient descent *∂F*/*∂***s**_*τ*_ = 0 and *∂F*/*∂***θ** = *0*, the update rules are derived as follows:

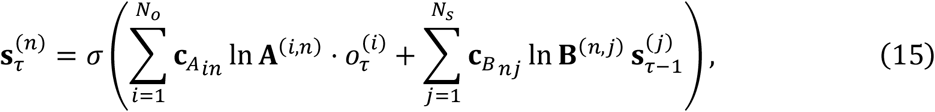

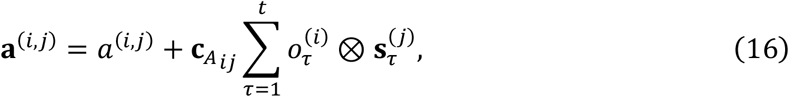

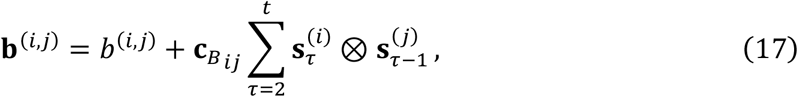

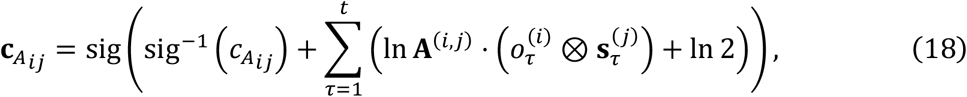

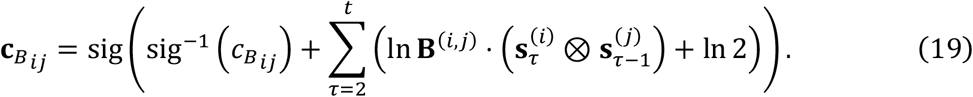

Equations (18) and (19) are equivalent to the equations for the dynamics of *C*_*W*_ and *C*_*K*_ in the neural network formation that are derived from the fixed point of the gradient descent on *L, ∂L*/*∂C*_*W*_ = *0* and *∂L*/*∂C*_*K*_ = *0*. That is, Eqs. (18) and (19) are plausible synaptic pruning rules for the canonical neural network.

### 4.3 Properties of connectivity parameters

In this subsection, we show that 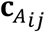 converges to 0 when 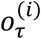 is randomly determined and independent of all elements of *s*_τ_, and the likelihood mapping posterior **A**^(i,j)^ and the mean state posterior 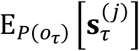 are sufficiently flat. The direction of the change in 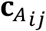 is determined by the sign of the second term on the right-hand side of Eq. (18). We adopt the following approximation for analytical tractability: In 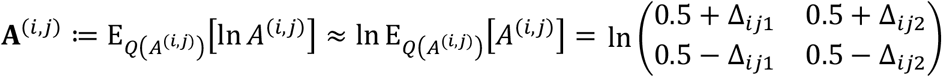. This approximation holds when **a**^(i,j)^ is sufficiently large, owing to the properties of the digamma functions (Isomura & Friston, 2020). Subsequently, the Taylor expansion of ln **A**^(i,j)^ at 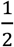 yields the following equation:

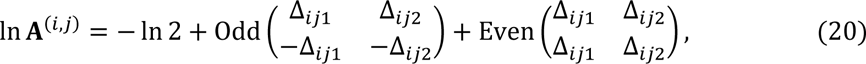

where Odd(*x*) and Even(*x*) are odd and even function components of ln(0.5 + *x*). This equation allows the following transformation:

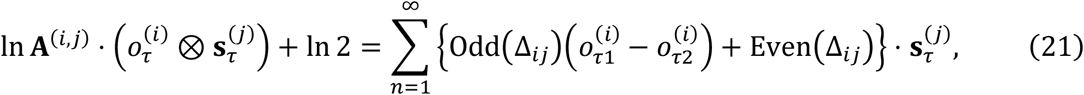

where Δ_*ij*_ and 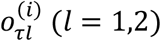 are given by Δ_*ij*_ ≔ (Δ_*ij1*_, Δ_*ij*_)^T^ and 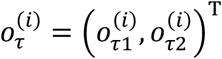. Because we consider the case where 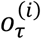 is independent of all elements of *S_τ_*, 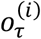 is independent of all the other observation. Here and after, 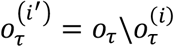 denotes the set of all the observations other than *i*th observation 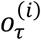. The expectation of In 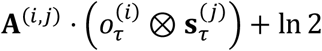 over observations is as follows:

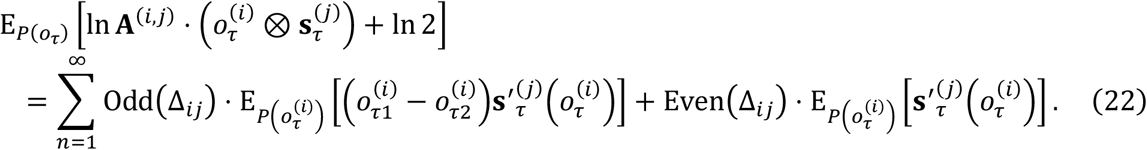

where 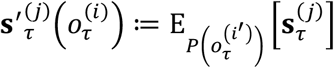 denotes the expected state posterior inferred without observing 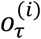. This is a random variable that depends on 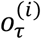. Because we consider the case in which 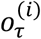 changes randomly, i.e., 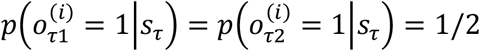, the expectation of 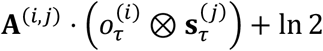 over observations is calculated as follows:

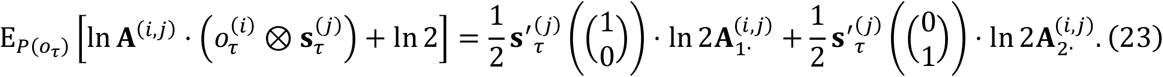

Furthermore, in the case where | Δ*ij* | ≪ 1 holds, In 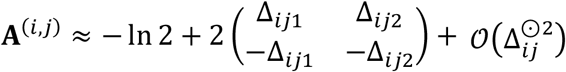 holds. With this approximation, the Taylor expansion of 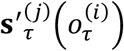 provides the following approximation:

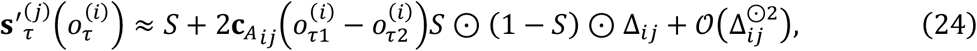

where *S* is defined by 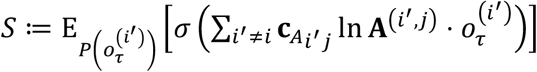. Besides, 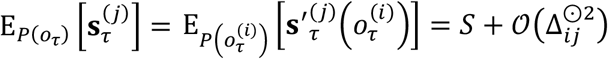 holds in this case. Subsequently, when 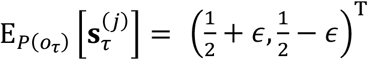 with |*ϵ*| *≪* 1 holds, the expectation of interest is approximated as follows:

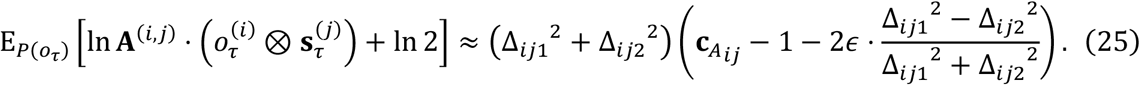

This means that the expectation above is negative when 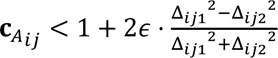 holds, that is, when 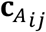 is not sufficiently close to 1. In addition, 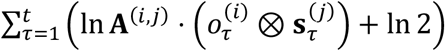 has a 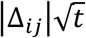-order standard deviation according to the central limit theorem. Taken together,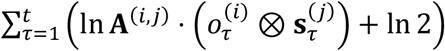 is a |Δ_*ij*_|^2^ *t* -order negative value with a 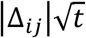- order standard deviation, which decreases monotonically with time when *t* is sufficiently large. Therefore, the direction of change in 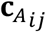 is negative when 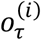 randomly determined and independent of all elements of *s*_τ_, and **A**^(i,j)^ and 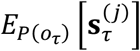 are sufficiently flat. Hence, 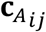 converges to 0 after sufficient sessions.

### 4.4 Numerical simulation 1: Blind source separation

The agents received 1024 (i.e., 32 × 32 pixels) observations that were generated using the generative model defined in Eq. (2). The model comprised two states, *s*^(1)^ and *s*^(2)^, which generated the observations. Both states randomly took values of 0 or 1 with a probability of 0.5 each (as depicted by the flat matrix *B* with all elements of 0.5). When *s*^(1)^ or *s*^(2)^ was ON, the shape of a circle or rectangle was displayed on the screen as sensory input for the agent (Fig. 2a). This was implemented using the following matrix *A*:

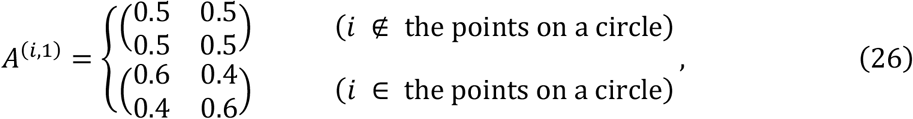

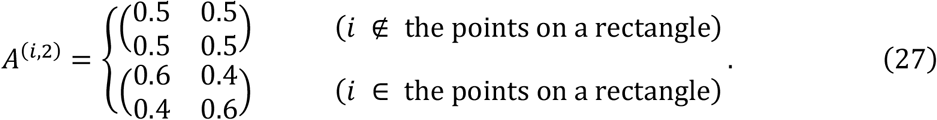

The agent optimized **s, a**, and **c**_**A**_ (i.e., **b** and **c**_**B**_ were omitted from this simulation). The prior beliefs about *A* and *C*_*A*_ characterized the inference of the agents. The elements of prior belief *a*^(i,j)^ with the *i* -index denoting a point on the shape is expressed as follows: 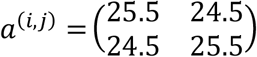. The other elements *a*^(i,j)^ were randomly sampled from a uniform distribution with the center of 25 and interval of 1. All elements of prior belief about *C*_*A*_ were sampled from a uniform distribution with the center of 0.9 and interval of 0.1. Further details are provided in the source code.

The full model and BSyMP agents optimized their posterior beliefs: they optimized **s**_τ_ at every time point and optimized **a** and **c**_**A**_ at every 10 time points. Following optimization over 10,000 time points, BMR was applied to the learned full model (Friston et al., 2019). BMR computed the change in free energy Δ*F*^(i,j)^ when each *a*^(i,j)^ was varied. A prior for reduction ã^(i,j)^ was set to a uniform matrix with large positive values. The element of the posterior **a**^(i,j)^ was reduced if the change in the approximate free energy Δ*F*^(i,j)^ ≔ ln ℬ(ã ^(i,j)^) − ln ℬ (**ã**^(i,j)^) − ln ℬ(*a*^(i,j)^) + ln ℬ(**a**^(i,j)^) was negative and retained otherwise. Further details regarding BMR can be found in Friston et al., 2019.

After performing BMR, the variational free energy was calculated by following Eq. (4) at the end of the training sessions, however, the complexity of the connectivity parameters was excluded for a consistent evaluation among the three methods. Subsequently, the normalized mapping posteriors for all agents were calculated. The normalized mapping posterior **A** for the full model and BMR and 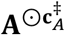 for BSyMP are expressed as follows:

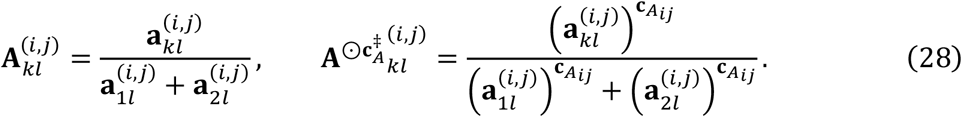

Finally, 1000 time-point-length data points were created using the same generative process to test the performance of the agents.

### 4.5 Numerical simulation 2: Rule learning

The agents received 3072 (i.e., 32 × 32 × 3 pixels) observations generated using the generative model defined in Eq. (2). The model comprised 12 states, each of which exhibited a certain shape on one of the three screens. Matrix *A* was set similarly to that in Simulation 1, but the fidelity of the observations was determined as follows:

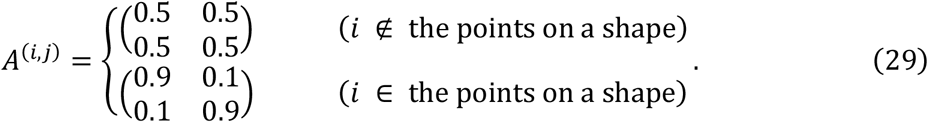

The state change followed the transition rule characterized by matrix *B*. All 2 × 2 submatrices were symmetric. Therefore, 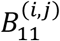 was sufficient to characterize *B*^(i,j)^, as follows:

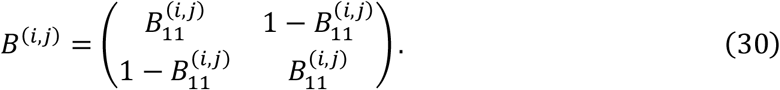

The initial states were determined randomly, as depicted by a flat block vector *D* with all elements of 0.5.

The agent employed the prior beliefs about *A, B* and *C*_*A*_, *C*_*B*_. The elements of prior belief *a*^(i,j)^ were set as follows:

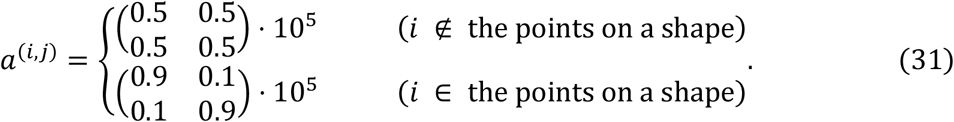

All elements of the prior belief *b*^(i,j)^ were sampled from a uniform distribution with the mean of 50 and interval of 2. All elements of the prior beliefs *c*_*A*_ and *c*_*B*_ were sampled from a uniform distribution with the mean of 0.9 and interval of 0.1. Further details are provided in the source code.

The full model and BSyMP agents optimized their posterior beliefs: they optimized **s**_*r*_ at every time point and optimized **a, b, c**_*A*_, and **c**_*B*_ at every 100 time points. Following optimization over 10,000 time points, BMR was applied to the learned full model in the same manner as in Simulation 1.

After performing BMR, the variational free energy and the normalized transition rule posteriors **B** and 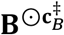 were calculated in the same manner as in Simulation 1. Finally, 1000 time point-length data points were created using the same generative process to test the prediction performance of the agents. The predicted state of the *i*-th source was defined as 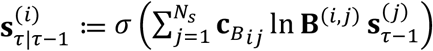 for BSyMP and 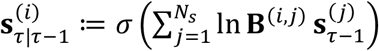 for the full model and BMR. The actual probability of the *i*-th source state was defined as follows: 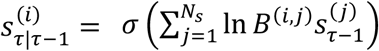.

## Acknowledgment

T.I. is funded by the Japan Society for the Promotion of Science (JSPS) KAKENHI (Ref: JP23H04973) and the Japan Science and Technology Agency (JST) CREST (Ref: JPMJCR22P1). The funders had no role in study design, data collection and analysis, decision to publish, or preparation of the manuscript.

## CRediT authorship contribution statement

Ukyo T. Tazawa: Conceptualization, Data curation, Formal analysis, Investigation, Methodology, Software, Visualization, Writing – original draft, Writing – review & editing.

Takuya Isomura: Conceptualization, Formal analysis, Funding acquisition, Methodology, Project administration, Resources, Supervision, Writing – review & editing.

