## Appendix for "Synaptic pruning facilitates online Bayesian model selection"

### Appendix A. Performance of online version of BMR on the BSS task

As noted in section 2.2, the underperformance of BMR in the BSS task can be attributed to the post hoc model reduction. Here we evaluated the online version of BMR (Friston et al., 2017). The online BMR agent performed BMR on each session and continued inference and learning with the reduced posterior. This indicates that once a parameter was reduced in a session, it remained reduced in the subsequent sessions. Thus, to reduce only the parameters with high confidences for the reduction, the element $\mathbf{a}^{(i,j)}$ was reduced when $\Delta F^{(i,j)} < -3$ held (see Section 4.4 for further detail).

The performance of the online BMR agent was almost the same level as that of the BMR agent (Figs. A.1). This is arguably because the careful reduction led to small effect on the mapping posterior at the early stage of learning, resulting in the failure in modifying the learning trajectories from those of the full model. We also evaluated online BMR that reduced $\mathbf{a}^{(i,j)}$ when $\Delta F^{(i,j)} < 0$ held—the same threshold for the post hoc version—almost all the parameters were reduced in this case, resulting in the performance close to the chance level (data not shown). In summary, online BMR as well as conventional BMR failed to perform the BSS task because of the heuristic in the threshold for $\Delta F^{(i,j)}$.

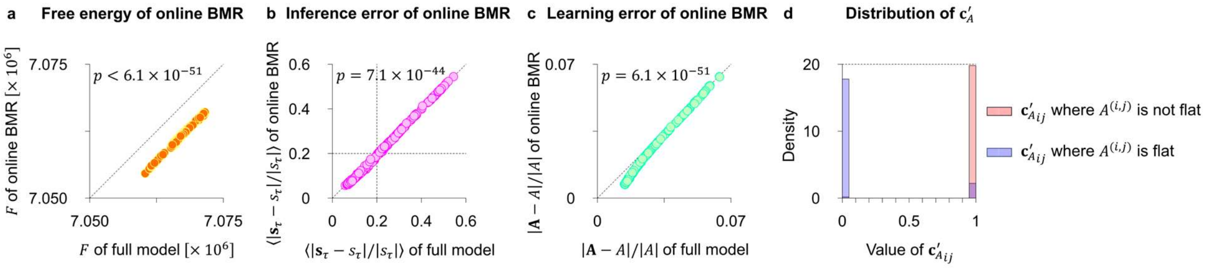


**Fig. A.1. Performance of online version of BMR on the BSS task.** (**a**) A comparison of the free energy between online BMR and the full model. (**b**) A comparison of the state inference error between online BMR and the full model. (**c**) A comparison of the parameter learning error between online BMR and the full model. (**d**) The distribution of the online BMR's counterpart, $\mathbf{c}'_A$, to the connectivity parameter in BSyMP, $\mathbf{c}_A$, at the end of the training sessions.

**Appendix B. Analysis on the reduction by BMR**

Here we analyze the mechanism of BMR reducing an element of A matrix. BMR reduces the element if the change in free energy $\Delta F^{(i,j)} \coloneqq F_{\mathrm{red}(i,j)} - F_{\mathrm{full}}$ is negative, where $F_{\mathrm{full}}$ and $F_{\mathrm{red}(i,j)}$ are free energies of full model and that with the reduced $(i,j)$ connectivity, respectively. Under the POMDP generative model, this is computed as $\Delta F^{(i,j)} = \ln \mathcal{B}\big(\tilde{a}^{(i,j)}\big) - \ln \mathcal{B}\big(\tilde{\mathbf{a}}^{(i,j)}\big) - \ln \mathcal{B}\big(a^{(i,j)}\big) + \ln \mathcal{B}\big(\mathbf{a}^{(i,j)}\big) = \sum_{l=1}^{2} \left\{ \ln \mathcal{B}\big(\tilde{a}_{\cdot l}{}^{(i,j)}\big) - \ln \mathcal{B}\big(\tilde{\mathbf{a}}_{\cdot l}{}^{(i,j)}\big) - \ln \mathcal{B}\left(a_{\cdot l}^{(i,j)}\right) + \ln \mathcal{B}\left(\mathbf{a}_{\cdot l}^{(i,j)}\right) \right\}$ . We analyze the behavior of $\Delta F_{\cdot l}^{(i,j)} \coloneqq \ln \mathcal{B}\big(\tilde{a}_{\cdot l}{}^{(i,j)}\big) - \ln \mathcal{B}\big(\tilde{\mathbf{a}}_{\cdot l}{}^{(i,j)}\big) - \ln \mathcal{B}\left(a_{\cdot l}^{(i,j)}\right) + \ln \mathcal{B}\left(\mathbf{a}_{\cdot l}^{(i,j)}\right)$ under the following conditions:

$$a_{\cdot l}^{(i,j)} = \begin{pmatrix} a \\ a \end{pmatrix}, \qquad \mathbf{a}_{\cdot l}^{(i,j)} = m \begin{pmatrix} a + d \\ a - d \end{pmatrix}, \qquad \tilde{a}_{\cdot l}{}^{(i,j)} = \begin{pmatrix} \tilde{a} \\ \tilde{a} \end{pmatrix},$$

$$1 \le m, \qquad -\frac{m-1}{m} a \le d \le \frac{m-1}{m} a, \qquad \tilde{a} \gg ma. \tag{B.1}$$

These conditions enable us to handle $\mathbf{a}_{\cdot l}^{(i,j)}$ as a function of $m$ and $d$, and interpret the meaning of the posterior easily. Coefficient $m$ increases as the learning proceeds, and bias $d$ denotes how $\mathbf{a}_{\cdot l}^{(i,j)}$ is far from a flat posterior. The ranges of $m$ and $d$ are determined to assure $a_{kl}^{(i,j)} \le \mathbf{a}_{kl}^{(i,j)}$. With these notations, $\Delta F_{\cdot l}^{(i,j)}$ is calculated as follows:

$$\Delta F_{\cdot l}^{(i,j)} \approx m\{(a+d)\ln(a+d) + (a-d)\ln(a-d) - 2a\ln a\} + \frac{1}{2}\left(\ln\frac{a}{a+d} + \ln\frac{a}{a-d} - \ln m\right). \tag{B.2}$$

This calculation was based on Stirling's approximation: $\ln \Gamma(x) \approx x \ln x - x - \frac{1}{2} \ln x$ . Taking derivative of $\Delta F_{\cdot l}^{(i,j)}$ regarding $d$ tells us that $\Delta F_{\cdot l}^{(i,j)}$ is a convex function of $d$ and

$\underset{d}{\operatorname{argmin}}\, \Delta F_{\cdot l}^{(i,j)} = 0$ gives the minimum value in the range of Eq. (B.1). Thus, $\min_{d} \Delta F_{\cdot l}^{(i,j)} = -\frac{1}{2}\ln m \,(\leq 0)$. When $d$ is close to 0, the following approximation holds:

$$\Delta F_{\cdot l}^{(i,j)} \approx \Delta F_{\cdot l}^{(i,j)}\Big|_{d=0} + \left(\frac{\partial \Delta F_{\cdot l}^{(i,j)}}{\partial d}\Bigg|_{d=0}\right) d + \left(\frac{\partial^2 \Delta F_{\cdot l}^{(i,j)}}{\partial d^2}\Bigg|_{d=0}\right) d^2 = -\frac{1}{2}\ln m + \frac{2(ma+1)}{a^2} d^2. \quad (\mathrm{B.3})$$

Therefore, the solution of $\Delta F_{\cdot l}^{(i,j)} = 0$ is approximated as $d = \pm\frac{2}{a}\sqrt{\frac{\ln m}{ma+1}}$, indicating that the larger $m$ becomes, the narrower the range of $d$ that holds $\Delta F_{\cdot l}^{(i,j)} < 0$ becomes. In summary, the range of BMR reducing sub-matrix to flat becomes narrower monotonically as the learning proceeds. This behavior is equivalent to the weight-based rule that prunes synapses with small weights in neural networks.

### Appendix C. Correspondence between BSyMP and BMR

Here we show the correspondence between BSyMP and the previously developed Bayesian model reduction (BMR) method. BMR compares the amount of free energy between the learned full model and reduced model with different priors. For the factorial PODMP model considered in this work, the reduction of an element of *A* matrix is achieved by replacing its flat prior $a^{(i,j)} = \begin{pmatrix} \alpha & \alpha \\ \alpha & \alpha \end{pmatrix}$ with strong flat prior $\tilde{a}^{(i,j)} = \begin{pmatrix} \tilde{\alpha} & \tilde{\alpha} \\ \tilde{\alpha} & \tilde{\alpha} \end{pmatrix}$, where $0 < \alpha \ll \tilde{\alpha}$. The element of the reduced posterior $\tilde{\mathbf{a}}^{(i,j)}$ takes the form of $\tilde{\mathbf{a}}^{(i,j)} = \tilde{a}^{(i,j)} + \sum_{\tau=1}^{t} o_\tau^{(i)} \otimes \mathbf{s}_\tau^{(j)}$. Thus, the normalized reduced posterior $\widetilde{\mathbf{A}}_{kl}^{(i,j)}$ is provided as follows:

$$\widetilde{\mathbf{A}}_{kl}^{(i,j)} := \frac{\tilde{\mathbf{a}}_{kl}^{(i,j)}}{\tilde{\mathbf{a}}_{1l}^{(i,j)} + \tilde{\mathbf{a}}_{2l}^{(i,j)}} = \frac{\tilde{\alpha} + \sum_{\tau=1}^{t} o_{\tau k}^{(i)} \mathbf{s}_{\tau l}^{(j)}}{\tilde{\alpha} + \sum_{\tau=1}^{t} o_{\tau 1}^{(i)} \mathbf{s}_{\tau l}^{(j)} + \tilde{\alpha} + \sum_{\tau=1}^{t} o_{\tau 2}^{(i)} \mathbf{s}_{\tau l}^{(j)}} = \frac{\tilde{\alpha} + \sum_{\tau=1}^{t} o_{\tau k}^{(i)} \mathbf{s}_{\tau l}^{(j)}}{2\tilde{\alpha} + \sum_{\tau=1}^{t} \mathbf{s}_{\tau l}^{(j)}}. \quad (\mathrm{A.1})$$

Conversely, the posterior belief obtained by BSyMP ($\mathbf{A}^{\mathbf{c}_A}{}_{kl}^{(i,j)}$) takes the following form:

$$\mathbf{A}^{\mathbf{c}_A}{}_{kl}^{(i,j)} := \frac{\left(\alpha + \sum_{\tau=1}^{t} o_{\tau k}^{(i)} \mathbf{s}_{\tau l}^{(j)}\right)^{\mathbf{c}_{Aij}}}{\left(\alpha + \sum_{\tau=1}^{t} o_{\tau 1}^{(i)} \mathbf{s}_{\tau l}^{(j)}\right)^{\mathbf{c}_{Aij}} + \left(\alpha + \sum_{\tau=1}^{t} o_{\tau 2}^{(i)} \mathbf{s}_{\tau l}^{(j)}\right)^{\mathbf{c}_{Aij}}}. \quad (\mathrm{A.2})$$

Because we consider the case with a flat prior $a^{(i,j)} = \begin{pmatrix} \alpha & \alpha \\ \alpha & \alpha \end{pmatrix}$, $\mathbf{A}^{\mathbf{c}_A}{}_{kl}^{(i,j)}$ can be transformed as follows:

$$\mathbf{A}^{\mathbf{c}_A}{}_{kl}^{(i,j)} = \frac{\left(1 + \frac{2\left(\alpha + \sum_{\tau=1}^{t} o_{\tau k}^{(i)} \mathbf{s}_{\tau l}^{(j)}\right)}{\sum_{\tau=1}^{t} \mathbf{s}_{\tau l}^{(j)}} - 1\right)^{\mathbf{c}_{Aij}}}{\left(1 + \frac{2\left(\alpha + \sum_{\tau=1}^{t} o_{\tau 1}^{(i)} \mathbf{s}_{\tau l}^{(j)}\right)}{\sum_{\tau=1}^{t} \mathbf{s}_{\tau l}^{(j)}} - 1\right)^{\mathbf{c}_{Aij}} + \left(1 + \frac{2\left(\alpha + \sum_{\tau=1}^{t} o_{\tau 2}^{(i)} \mathbf{s}_{\tau l}^{(j)}\right)}{\sum_{\tau=1}^{t} \mathbf{s}_{\tau l}^{(j)}} - 1\right)^{\mathbf{c}_{Aij}}}. \quad (\mathrm{A.3})$$

As the Taylor expansion of $(1+x)^c$ at $x=0$ yields $(1+x)^c = 1 + cx + \mathcal{O}(x^2) \approx 1 + cx$ for small $x$, we obtain the following approximation:

$$
\begin{aligned}
\mathbf{A}^{\mathbf{c}_A}{}_{kl}^{(i,j)} &\approx \frac{1 + \left(\dfrac{2\left(\alpha + \sum_{\tau=1}^{t} o_{\tau k}^{(i)} \mathbf{s}_{\tau l}^{(j)}\right)}{\sum_{\tau=1}^{t} \mathbf{s}_{\tau l}^{(j)}} - 1\right)\mathbf{c}_{A_{ij}}}{1 + \left(\dfrac{2\left(\alpha + \sum_{\tau=1}^{t} o_{\tau 1}^{(i)} \mathbf{s}_{\tau l}^{(j)}\right)}{\sum_{\tau=1}^{t} \mathbf{s}_{\tau l}^{(j)}} - 1\right)\mathbf{c}_{A_{ij}} + 1 + \left(\dfrac{2\left(\alpha + \sum_{\tau=1}^{t} o_{\tau 2}^{(i)} \mathbf{s}_{\tau l}^{(j)}\right)}{\sum_{\tau=1}^{t} \mathbf{s}_{\tau l}^{(j)}} - 1\right)\mathbf{c}_{A_{ij}}} \\
&= \frac{\alpha + \frac{1}{2}\left(\dfrac{1}{\mathbf{c}_{A_{ij}}} - 1\right)\sum_{\tau=1}^{t} \mathbf{s}_{\tau l}^{(j)} + \sum_{\tau=1}^{t} o_{\tau k}^{(i)} \mathbf{s}_{\tau l}^{(j)}}{2\left\{\alpha + \frac{1}{2}\left(\dfrac{1}{\mathbf{c}_{A_{ij}}} - 1\right)\sum_{\tau=1}^{t} \mathbf{s}_{\tau l}^{(j)}\right\} + \sum_{\tau=1}^{t} \mathbf{s}_{\tau l}^{(j)}}.
\end{aligned} \tag{A.4}
$$

Therefore, Eq. (A.4) is homologous to Eq. (A.1) when setting $\tilde{\alpha} = \alpha + \frac{1}{2}\left(\frac{1}{\mathbf{c}_{A_{ij}}} - 1\right)\sum_{\tau=1}^{t} \mathbf{s}_{\tau l}^{(j)}$, indicating the formal correspondence between BSyMP and BMR. This analysis further highlights an important contrast between BSyMP and BMR. BMR is a binary reduction method that reduces an element if the reduction decreases free energy since $\tilde{\alpha}$ is heuristically set as a large value a priori. Conversely, BSyMP is a continuous reduction method that reduces an element depending on the value of $\alpha + \frac{1}{2}\left(\frac{1}{\mathbf{c}_{A_{ij}}} - 1\right)\sum_{\tau=1}^{t} \mathbf{s}_{\tau l}^{(j)}$, which provides the Bayes-optimal solution for $\mathbf{A}^{\mathbf{c}_A}{}_{kl}^{(i,j)}$.

### Appendix D. Approximation on partition functions

Here, we derive the approximation of partition function: $E_Q\left[\ln Z_{C_A}\right] \approx \ln 2 \cdot \sum_{i=1}^{N_o}\left(1 - \sum_{j=1}^{N_s} \mathbf{c}_{A_{ij}}\right)$. When $A^{(i,j)}$ is expressed as $A^{(i,j)} = \begin{pmatrix} 0.5+\Delta_{ij1} & 0.5+\Delta_{ij2} \\ 0.5-\Delta_{ij1} & 0.5-\Delta_{ij2} \end{pmatrix}$, the following equation holds:

$$Z_{C_A}^{(i)} := \prod_{j=1}^{N_s} A_{1\cdot}^{(i,j)^{C_{Aij}}} s_\tau^{(j)} + \prod_{j=1}^{N_s} A_{2\cdot}^{(i,j)^{C_{Aij}}} s_\tau^{(j)}$$

$$= \prod_{j=1}^{N_s} \left((0.5, 0.5) + \Delta_{ij\cdot}\right)^{C_{Aij}} s_\tau^{(j)} + \prod_{j=1}^{N_s} \left((0.5, 0.5) - \Delta_{ij\cdot}\right)^{C_{Aij}} s_\tau^{(j)}, \qquad (\mathrm{C}.1)$$

where $\Delta_{ij\cdot} := \left(\Delta_{ij}\ , \Delta_{ij2}\right)$. Then we obtain the following approximation by adopting the first order approximation on Δ.

$$Z_{C_A}^{(i)} \approx \prod_{j=1}^{N_s} (0.5, 0.5)^{C_{Aij}} s_\tau^{(j)} \left\{ 1 + \frac{\Delta_{i1\cdot}^{C_{Ai1}} s_\tau^{(1)}}{\left((0.5, 0.5)^{C_{Ai1}}\right) s_\tau^{(1)}} + \cdots + \frac{\Delta_{iN_s\cdot}^{C_{AiN_s}} s_\tau^{(N_s)}}{\left((0.5, 0.5)^{C_{AiN_s}}\right) s_\tau^{(N_s)}} \right\}$$

$$+ \prod_{j=1}^{N_s} (0.5, 0.5)^{C_{Aij}} s_\tau^{(j)} \left\{ 1 - \frac{\Delta_{i1\cdot}^{C_{Ai1}} s_\tau^{(1)}}{\left((0.5, 0.5)^{C_{Ai1}}\right) s_\tau^{(1)}} - \cdots - \frac{\Delta_{iN_s\cdot}^{C_{AiN_s}} s_\tau^{(N_s)}}{\left((0.5, 0.5)^{C_{AiN_s}}\right) s_\tau^{(N_s)}} \right\}$$

$$= 2 \prod_{j=1}^{N_s} \left((0.5\ 0.5)^{C_{Aij}}\right) s_\tau^{(j)} = 2 \prod_{j=1}^{N_s} 0.5^{C_{Aij}}. \qquad (\mathrm{C}.2)$$

The last equality holds because $\sum_{l=1}^{2} s_{\tau l}^{(j)} = 1$ holds. With this approximation, $E_Q\left[\ln Z_{C_A}^{(i)}\right]$ is calculated as follows:

$$E_Q\left[\ln Z_{C_A}^{(i)}\right] \approx \ln 2 + E_Q\left[\sum_{j=1}^{N_s} 0.5^{C_{A_{ij}}}\right] = \ln 2 + \sum_{j=1}^{N_s}\left(\mathbf{c}_{A_{ij}} \ln 0.5 + \left(1 - \mathbf{c}_{A_{ij}}\right)\ln 1\right) = \ln 2 \cdot \left(1 - \sum_{j=1}^{N_s} \mathbf{c}_{A_{ij}}\right). \quad (\mathrm{C}.3)$$

Therefore, $E_Q\left[\ln Z_{C_A}\right] = \mathrm{E}_Q\left[\sum_{i=1}^{N_o} \ln Z_{C_A}^{(i)}\right] \approx \ln 2 \cdot \sum_{i=1}^{N_o}\left(1 - \sum_{j=1}^{N_s} \mathbf{c}_{A_{ij}}\right)$. In the same manner, $E_Q\left[\ln Z_{B^{c_{B_{s_{\tau-1}}}}}\right] \approx \ln 2 \cdot \sum_{i=1}^{N_s}\left(1 - \sum_{j=1}^{N_s} \mathbf{c}_{B_{ij}}\right)$ is derived.
